# Concordance of Race Documented in Electronic Health Records and Genetic Ancestry

**DOI:** 10.1101/598706

**Authors:** Theresa L. Walunas, Jennifer A. Pacheco, Kathryn L. Jackson, Anna Roberts, Loren L. Armstrong, Jess Behrens, M. Geoffrey Hayes, Abel N. Kho

## Abstract

**Objective:** Genetic screening is the gold standard for biogeographical ancestry (i.e. race), but this information is often unavailable to those developing research studies. We assessed agreement between census- and electronic health record (EHR)-derived demographic data with genetic ancestry to determine if these sources could support selection of diverse cohorts.

**Materials and Methods:** We identified a population of 4,837 genotyped patients and determined concordance between genetic measures of ancestry against race derived from decennial nationwide census, electronic medical records, and self-report.

**Results:** We identified a 90% or greater concordance between the EHR-derived data and genetic ancestry. Census data had a high concordance (97%) with genetic and self-reported data for patients of European ancestry but low concordance for patients of African ancestry (64%).

**Discussion and Conclusions:** The high concordance between EHR-derived race and genetic ancestry suggests that EHR-derived information could be an effective proxy for race when recruiting for diverse research cohorts.

## BACKGROUND AND SIGNIFICANCE

The goal of precision medicine is to develop personally-targeted approaches to health care, by including individual genetic, social and environmental components into care planning. As focus on precision medicine grows, it has become increasing critical to be able to develop diverse patient populations to participate in research. Many research studies that are focused on personalized medicine, including the Electronic Medical Records and Genomics Network (eMERGE)[1–3] and The All of Us Research Program (previously the Precision Medicine Initiative),[4] seek to develop diverse cohorts of patients. Given their access to patient medical records and longitudinal patient information, medical centers and other health care providing organizations are often central to the development of such research cohorts and must develop strategies to engage study participants who represent the U.S. population at large to ensure that all patients can receive effective personalized care. Representativeness is based on a variety of information, including basic demographics such as sex, age, race, ethnicity, socioeconomic status and geographic location, all of which can be captured in the electronic health record (EHR).

The HITECH Act was passed in 2009 to accelerate the adoption of EHR systems by the nation’s healthcare providers.[5] Within HITECH, the Meaningful Use (MU) program attempted to catalyze adoption and effective use of the technology by requiring healthcare providers to demonstrate high quality data collection, including basic demographics, in order to receive incentive payments and avoid payment reductions from Medicare.[6] In 2010, only 12% of hospitals[7] and 26% of ambulatory providers[8] were using EHR systems to capture patient visits. By 2015, 96% of hospitals[9] and 78% of ambulatory providers[10] had adopted EHRs. The first phase of MU required documentation of demographics (including race and ethnicity) for 50% of unique patients seen by a given health care provider and the second phase of MU (initiated in 2014) increased that requirement to 80% As of 2015, those ambulatory and inpatient providers who participated in MU documented basic demographics on 92-100% of their patients.[11] In fact, this measure of effective EHR use was so successful that the Centers for Medicare and Medicaid Services determined it “topped out” and will no longer include it in the specific measures for MU or the Promoting Interoperability (formerly Advancing Care Information) component of the newly legislated Quality Payment Program.[12]

Increase in EHR usage as well as availability of reasonably complete demographic data within the EHR suggests a mechanism by which health care organizations recruiting for precision medicine-focused research studies, such as the All of Us Research program, could ensure diverse enrollment. While sex, age and geographic location are relatively easy to determine and validate, less is understood about how race/ethnicity information documented in EHRs correlates with genetic data, which is considered the gold standard for assessing biogeographic ancestry, or “race”.[13] Previous studies have explored how race documented by observers correlated with ancestry and discovered a high concordance between observer-documented race and biogeographic ancestry.[14] In this study, we extended those explorations, using a large patient cohort to assess the concordance of race identification through census data estimates, self-reporting, and EHR demographic data collected as part of health care and research activities with genome-wide estimates of biogeographic ancestry, to determine the best mechanism to ensure reasonable race/ethnicity diversity in patient recruitment efforts for precision medicine-focused research study cohorts.

## MATERIALS AND METHODS

To perform this study, we identified a population of 4,837 patients within the Northwestern Medicine system (Chicago, IL) who had consented to genotyping and use of their EHR data for genomic research as part of the NUgene project.[15] From each patient, data on race or biogeographic ancestry was collected from four sources: the NUgene patient enrollment questionnaire, a geocoded home address matched to US Census 2010 data, EHR data extracted from the Northwestern Medicine Enterprise Data Warehouse (NMEDW) and biogeographic ancestry data gathered as part of genome-wide association studies using either the Illumina 660W or 1M BeadChip Array.

We estimated proportions of continental (European, African, and Asian) ancestry for the NUgene study participants using STRUCTURE v.2.3.3[16] under an admixture model of k=3 clusters with HapMap samples of known continental ancestry.[17 18] Plurality of proportion of continental ancestry was used to assign a single continental ancestry: The Yoruba from Ibadan, Nigeria (YRI) as African samples; the Japanese from Tokyo, Japan (JPT) and Han Chinese from Beijing, China (CHB) as Asian samples; and CEPH [Utah residents with ancestry from northern and western Europe (CEU)] as European samples. We identified NUgene participants as “Black or African American” if they had 25% (equivalent to one of four grandparents) or more YRI ancestry, which approximately reflects the point of transition along the sigmoidal curve of ancestry from European to African ancestry (see Figure 1), and may reflect the impact of historical practices in race identification in the U.S. such as the one-drop rule which considered a mixed race person with any black ancestor as black.[19] Additional studies comparing self-reported race and genetic ancestry in African and African American cohorts divided the percentage of African ancestry into quartiles and found that no one with 25% YRI or less identified themselves as having African ancestry.[20–22] Conversely, we assigned “White or European American” to those with 75% or more CEU. The few study subjects not meeting either of these criteria were instead assigned “Other” as done for the assignments from census data.

**Figure 1:**
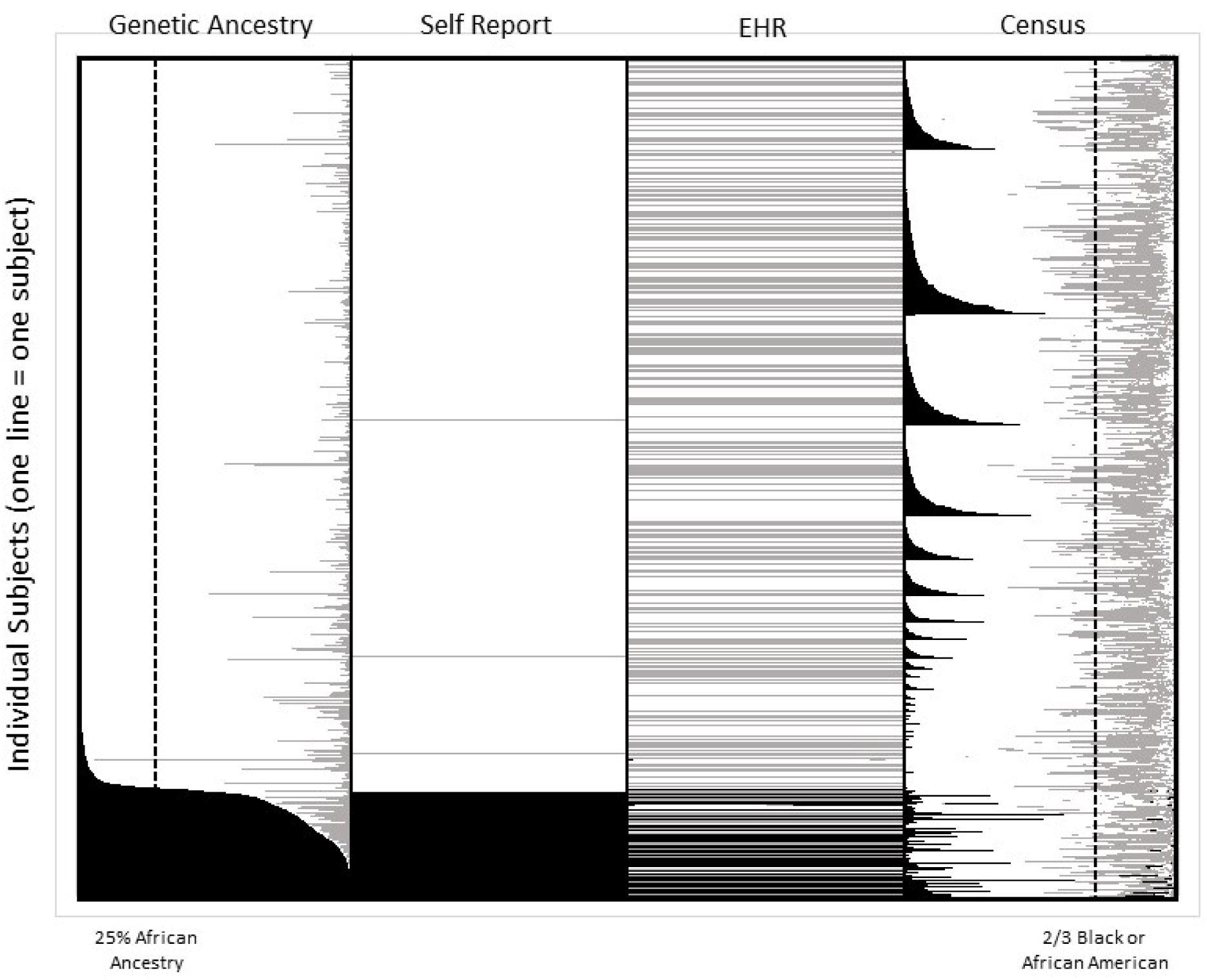
Classification of race based on genetic ancestry, self-report, EHR and census data. Each horizontal line represents a single individual’s data. For genetic ancestry, black represents the estimated proportion of African ancestry, white represents the estimated proportion of European ancestry, and grey represents the proportion of Asian ancestry. For self-report and EHR data, black represents Black or African American, white represents White/Caucasian, and grey represents Other. For census data, black represents the proportion of Black or African American residents in the study subject’s census tract, white is the proportion of white residents in the study subject’s census tract and grey is the proportion of other residents in the study subject’s census tract. The dashed lines illustrate cutoffs for genetic and census data. The plot is limited to those subjects with no missing data among the four sources and is sorted from top to bottom from low to high estimated proportion of African Ancestry.

To estimate race using census data, we divided the total number of residents pertaining to each race, Black and White, in each census tract by the total number of residents in each tract. Participants located within a census tract with a white population greater than two-thirds were assigned “White”, and those in tracts with a Black or African American population greater than two-thirds were assigned “Black”. For subjects not falling within those constraints we assigned category “Other”.

Once we had extracted (in the case of the EHR), inferred (census) or empirically estimated (genetic analysis) race data from each alternate source and structured it in a format consistent with the self-reported data, we used frequency procedure generated tables in SAS 9.2 to measure the percentage of agreement in race assignment between each source with the others. A summary of our race assignment methods for each source of data is provided in Table 1.

**Table 1.** Criteria for Race Assignment for Individuals

| Data Source | Race Assigned | Criteria/cutoff for assignment |
| --- | --- | --- |
| Questionnaire | White | Patient response on questionnaire |
|  | Black | Patient response on questionnaire |
|  | Other | Not applicable |
| Census | White | Per 2010 US Census data, population White only divided by Total population for each census tract. If resulting percentage was greater than 2/3, race White was assigned. |
|  | Black or African-American | Per 2010 US Census data, population Black or African American only divided by Total population for each census tract. If resulting percentage was greater than 2/3, race Black was assigned. |
|  | Other | Per US Census data, if neither White nor Black had a 2/3 majority in a tract, race Other was assigned. |
| EHR | White | Race White recorded in EHR by provider or staff |
|  | Black | Race Black recorded in EHR by provider or staff |
|  | Other | Race recorded in EHR by provider or staff besides White or Black (Asian, Other, Declined, null) |
| Genetic Analysis | European ancestry (White) | Using ancestry estimates of percentage CEU, YRI, and CHB-JPT from STRUCTURE analysis, White race assigned for percent CEU greater than or equal to .75 |
|  | African ancestry (black) | Using ancestry estimates of percentage CEU, YRI, and CHB-JPT from STRUCTURE analysis, Black race assigned for percent YRI greater than .25. |
|  | Other Ancestry | For those not falling under the criteria for White or Black, race Other was assigned. |

## RESULTS

In this study, we identified a broad range in the accuracy of proxies for biogeographical ancestry of race compared against an empirical estimate from genome-wide SNP data. Figure 1 and Table 2 show the concordance between genetic and self-reported race / biogeographic ancestry data with census-based and EHR-derived race. As reported previously,[22 23] a study subject’s self-reported information is in strong agreement with their genetic ancestry estimates (> 97% concordance within both Black / African ancestry and White / European ancestry patients) and serves as an excellent proxy.

**Table 2:** Concordance between Genetic, Self-Report, EHR-based and Census-based determination of Race

|  |  | Genetic |  |  | Self-Report |  |  |
| --- | --- | --- | --- | --- | --- | --- | --- |
|  |  | African Ancestry | European Ancestry | All | Black | White | All |
| Genetic | African Ancestry |  |  |  | 97.3% |  |  |
|  | European Ancestry |  |  |  |  | 98.7% |  |
|  | All |  |  |  |  |  | 98.4% |
| Self-Report | Black | 95.0% |  |  |  |  |  |
|  | White |  | 99.9% |  |  |  |  |
|  | All |  |  | 99.2% |  |  |  |
| EHR | Black | 90.2% |  |  | 91.2% |  |  |
|  | White |  | 92.3% |  |  | 92.3% |  |
|  | All |  |  | 92.0% |  |  | 92.1% |
| Census | Black | 63.5% |  |  | 62.3% |  |  |
|  | White |  | 97.0% |  |  | 97.1% |  |
|  | All |  |  | 92.6% |  |  | 92.5% |

However, self-reported information will often not be available before recruitment for those developing cohorts for precision medicine studies. We determined that concordance between EHR-derived data and genetic estimates of biogeographic ancestry is > 90% and > 91% between EHR-derived data and self-report. EHR-derived data is often evaluated for its level of completeness to determine its effectiveness as a tool for patient identification. In 2015, after successful participation in the MU, the percentage of patients in our genotyped population who were missing documented race in the EHR was only 13.5%, suggesting that in the absence of self-reported information, data from the EHR was of sufficient completeness to be used for identification of potential participants for diverse cohorts.

When we evaluated concordance between census-based estimates and biogeographic ancestry or self-reported race, it was high and comparable to EHR-reported data for White / European ancestry participants (97% in both cases) but much lower for Black / African ancestry participants (62-63%). This is not surprising given that census data is not specific to the individual.

## DISCUSSION AND CONCLUSIONS

In this study we examined the concordance of self-report, EHR-documented and geospatial determination of race compared to genetic ancestry and demonstrated that self-reported ancestry information and racial demographics documented in the EHR have high concordance (99% and 92%, respectively) and can be used as effective proxies for biogeographic ancestry in the absence of genetic information, while inference of ancestry through census information may be less effective for patients of non-white/non-European descent.

While genetic estimation of biogeographic ancestry has been possible for several decades, it is rare for any given individual to have this information in the EHR. Except for a small population of patients already in biobanks or participating in other studies associated with medical organizations that have developed cohorts for genetic research, this information is unlikely to be available for the development of recruitment strategies for the diverse cohorts of people that will be key to precision medicine focused research studies. A previous study, implemented before wide-spread EHR adoption, demonstrated a high concordance between observer reported-race and genetic ancestry.[14] Our study extends this work and compares EHR-documented race, self-reported race and census-derived race determination with biogeographic ancestry as estimated from genetic tests across the same population of patients. Census data had high concordance with ancestry for patients of European descent (97%), but was less reliable for patients of African descent (64%). That it is >60% for people of African descent may reflect the uniqueness of Chicago being one of the most diverse cities while also being one of the most segregated cities in the U.S..[24] Given this pattern, we would expect most other cities of the U.S. would have <60% concordance between census and genetic estimates of biogeographical ancestry, thus limiting the use of census tract information for selecting diverse patient populations for research even when address data is available. On the other hand, our data show a high concordance between EHR-derived race information, self-reported data and biogeographic ancestry as estimated from genetic tests. This finding of concordance between ancestry and EHR data, while encouraging, still leaves room for improvement in how race is collected in EHR systems. Although the Meaningful Use regulations encouraged direct collection of information from patients, some collection of this information is still derived from observation by a clinic registration clerk, and previous work has suggested that for multi-racial patients observation may not capture the nuances of self-reported data.[25 26] Thus, in the absence of self-report or genetic determination of ancestry, EHR-derived race information, especially now that federal initiatives have spurred an increased the level of completeness of this data in the EHR, provides an alternate option that could be used by healthcare organizations to help ensure recruitment of a racially diverse population for studies such as the All of Us Research Program that require racial diversity prior to patient reported or genetic screening.

## ACKNOWLEDGEMENTS

The authors would like to thank the NUgene participants.

## AUTHOR CONTRIBUTIONS

T.L.W. participated in the analysis, writing and review of the manuscript. J.A.P. collected data and participated in the analysis and review of the manuscript. K.L.J. participated in the analysis and review of the manuscript. A.R. participated in the data analysis and review of the manuscript. L.L.A. participated in the analysis of data for the manuscript. J.B. participated in the analysis of the census data and review of the manuscript. M.G.H. participated in the conceptualization, analysis and review of the manuscript. A.N.K participated in the conceptualization, analysis and review of the manuscript.

## DATA AND MATERIALS

The genetic data was deposited to dbGaP under study accession numbers phs000237.v1.p1 and phs000360.v1.p1

## REFERENCES

1. Gottesman O, Kuivaniemi H, Tromp G, et al. The Electronic Medical Records and Genomics (eMERGE) Network: past, present, and future. Genet. Med. 2013;15(10):761–71.

2. McCarty CA, Chisholm RL, Chute CG, et al. The eMERGE Network: a consortium of biorepositories linked to electronic medical records data for conducting genomic studies. BMC Med Genomics 2011;4:13|.

3. Kho AN, Pacheco JA, Peissig PL, et al. Electronic medical records for genetic research: results of the eMERGE consortium. Sci. Transl. Med. 2011;3(79):79|.

4. Collins FS, Varmus H. A new initiative on precision medicine. N Engl J Med 2015;372(9):793–5.

5. Blumenthal D. Launching HITECH. N Engl J Med 2010;362(5):382–5.

6. Blumenthal D, Tavenner M. The “meaningful use” regulation for electronic health records. N Engl J Med 2010;363(6):501–4.

7. DesRoches CM, Charles D, Furukawa MF, et al. Adoption of electronic health records grows rapidly, but fewer than half of US hospitals had at least a basic system in 2012. Health Aff (Millwood) 2013;32(8):1478–85.

8. Hsiao CJ, Jha AK, King J, Patel V, Furukawa MF, Mostashari F. Office-based physicians are responding to incentives and assistance by adopting and using electronic health records. Health Aff (Millwood) 2013;32(8):1470–7.

9. Office of the National Coordinator for Health Information Technology, Non-federal Acute Care Hospital Health IT Adoption and Use. Secondary Non-federal Acute Care Hospital Health IT Adoption and Use 2016. https://dashboard.healthit.gov/datadashboard/documentation/hospital-health-it-adoption-use-data-documentation.php.

10. Jamoom EW, Yang N. Table of Electronic Health Record Adoption and Use among Office-based Physicians in the U.S., by State. 2015 National Electronic Health Records Survey: National Center for Health Statistics, 2016.

11. Center for Medicare and Medicaid Services, Medicare and Medicaid EHR Incentive Programs 2014 Provider Attestation Data through March 20, 2015. https://www.cms.gov/Regulations-and-Guidance/Legislation/EHRIncentivePrograms/Downloads/AttestationPerformanceData_Feb2015.pdf.

12. Center for Medicare and Medicaid Services, Quality Payment Program: Promoting Interoperability Requirements. Secondary Quality Payment Program: Promoting Interoperability Requirements 2017. https://qpp.cms.gov/mips/promoting-interoperability.

13. Lee SS, Bolnick DA, Duster T, Ossorio P, Tallbear K. Genetics. The illusive gold standard in genetic ancestry testing. Science 2009;325(5936):38–9.

14. Dumitrescu L, Ritchie MD, Brown-Gentry K, et al. Assessing the accuracy of observer-reported ancestry in a biorepository linked to electronic medical records. Genet. Med. 2010;12(10):648–50.

15. Ormond KE, Cirino AL, Helenowski IB, Chisholm RL, Wolf WA. Assessing the understanding of biobank participants. Am J Med Genet A 2009;149A(2):188–98.

16. Hubisz MJ, Falush D, Stephens M, Pritchard JK. Inferring weak population structure with the assistance of sample group information. Mol Ecol Resour 2009;9(5):1322–32.

17. International HapMap C, Frazer KA, Ballinger DG, et al. A second generation human haplotype map of over 3.1 million SNPs. Nature 2007;449(7164):851–61.

18. International HapMap C. A haplotype map of the human genome. Nature 2005;437(7063):1299–320.

19. Omi M, Winant H. Racial Formation in the United States: From the 1960s to the 1990. New York, NY: Routledge, 1994.

20. Tang H, Peng J, Wang P, Risch NJ. Estimation of individual admixture: analytical and study design considerations. Genet Epidemiol 2005;28(4):289–301.

21. Jorgenson E, Tang H, Gadde M, et al. Ethnicity and human genetic linkage maps. Am J Hum Genet 2005;76(2):276–90.

22. Yaeger R, Avila-Bront A, Abdul K, et al. Comparing genetic ancestry and self-described race in African Americans born in the United States and in Africa. Cancer Epidemiol Biomarkers Prev 2008;17(6):1329–38.

23. Divers J, Redden DT, Rice KM, et al. Comparing self-reported ethnicity to genetic background measures in the context of the Multi-Ethnic Study of Atherosclerosis (MESA). BMC Genet 2011;12:28.

24. Williams A, Emamdjomeh A. America is more diverse thanever — but still segregated. Washington Post 2018 May 2, 2018.

25. Baker DW, Cameron KA, Feinglass J, et al. A system for rapidly and accurately collecting patients’ race and ethnicity. Am J Public Health 2006;96(3):532–7.

26. Hasnain-Wynia R, Baker DW. Obtaining data on patient race, ethnicity, and primary language in health care organizations: current challenges and proposed solutions. Health Serv. Res.2006;41(4 Pt 1):1501–18.

